# Frequent first-trimester pregnancy loss in rhesus macaques infected with African-lineage Zika virus

**DOI:** 10.1101/2022.12.09.519791

**Authors:** Jenna R. Rosinski, Lauren E. Raasch, Patrick Barros Tiburcio, Meghan E. Breitbach, Phoenix M. Shepherd, Keisuke Yamamoto, Elaina Razo, Nicholas P. Krabbe, Mason I. Bliss, Alexander D. Richardson, Morgan A. Einwalter, Andrea M. Weiler, Emily L. Sneed, Kerri B. Fuchs, Xiankun Zeng, Kevin K. Noguchi, Terry K. Morgan, Alexandra J. Alberts, Kathleen M. Antony, Sabrina Kabakov, Karla K. Ausderau, Ellie K. Bohm, Julia C. Pritchard, Rachel V. Spanton, James N. Ver Hoove, Charlene B. Y. Kim, T. Michael Nork, Alex W. Katz, Carol A. Rasmussen, Amy Hartman, Andres Mejia, Puja Basu, Heather A. Simmons, Jens C. Eickhoff, Thomas C. Friedrich, Matthew T. Aliota, Emma L. Mohr, Dawn M. Dudley, David H. O’Connor, Christina M. Newman

## Abstract

In the 2016 Zika virus (ZIKV) pandemic, a previously unrecognized risk of birth defects surfaced in babies whose mothers were infected with Asian-lineage ZIKV during pregnancy. Less is known about the impacts of gestational African-lineage ZIKV infections. Given high human immunodeficiency virus (HIV) burdens in regions where African-lineage ZIKV circulates, we evaluated whether pregnant rhesus macaques infected with simian immunodeficiency virus (SIV) have a higher risk of African-lineage ZIKV-associated birth defects. Remarkably, in both SIV+ and SIV-animals, ZIKV infection early in the first trimester caused a high incidence (78%) of spontaneous pregnancy loss within 20 days. These findings suggest a significant risk for early pregnancy loss associated with African-lineage ZIKV infection and provide the first consistent ZIKV-associated phenotype in macaques for testing medical countermeasures.

## Body Text

Zika virus (ZIKV) is a flavivirus discovered in 1947 in Uganda^1^. ZIKV was historically associated with intermittent epidemics throughout Africa, Asia, and Oceania, resulting in mild illness with seemingly few consequences. When the virus emerged in Brazil in 2015, there was an increase in cases of infant microcephaly^2^. This increase in microcephaly and other developmental abnormalities among neonates was ultimately associated with ZIKV exposure in-utero, drawing the attention of the broader scientific community to congenital Zika syndrome (CZS)^3– 5^. In the United States, 5-10% of infants with known gestational ZIKV exposure have developmental outcomes consistent with CZS^6^. Although the public health emergency has ended, recent outbreaks in India and evidence of periodic human infections elsewhere suggest ZIKV remains a threat during pregnancy^7–10.^.

Macaques have been used to model ZIKV infection during pregnancy using varying gestational time points, strains, doses, and routes of infection. Due to interest in the 2016 ZIKV pandemic, most studies have used Asian-lineage viruses, and in these studies, infection earlier in gestation frequently led to more severe outcomes^11–13^. Previously, we found a fetal demise rate of 26% (n=50) when using an Asian-lineage strain (PRVABC59) to infect macaques during the first trimester (<GD 55)^14^. Across multiple more recent studies using this strain, adverse fetal outcomes remain relatively rare (<10%; n=21) (Extended Data Table 1)^12,13,15–17^.

This study was designed to complement the NIH-supported International Prospective Cohort Study of HIV and Zika in Infants and Pregnancy (HIV ZIP)^18^. The goals of HIV ZIP are to determine whether infection with HIV and treatment with antiretroviral therapy (ART) increases the risk for ZIKV infection in the fetus and assess the risk of fetal co-infection with HIV and ZIKV. Unlike human co-infections, macaques can be infected with the same dose, strain, and route of ZIKV and simian immunodeficiency virus (SIV)^19^. SIV-induced disease in macaques manifests similarly to HIV-induced disease in humans, making SIV-infected macaques a useful model for the study of human HIV infection^20^. Areas of high HIV prevalence in sub-Saharan Africa also overlap with areas of historical ZIKV infections^21–23^. Additionally, African-lineage ZIKV strains show greater pathogenicity in mice than Asian-lineage strains and resulted in resorption of all embryos in dams infected with African-lineage ZIKV^24,25^. Therefore, we used an African-lineage virus from Senegal (ZIKV-DAK; strain 41524) in this study. This is the only ZIKV strain in this study; hereafter it is referred to as ZIKV. When we previously infected macaques with a high dose (1×10^8^ PFU) of this strain at gestational day (GD) 45, we observed a demise rate of 38% (n=8), whereas a physiological dose (1×10^4^ PFU) resulted in no demise (n=4)^26,27^.

For this study, twenty-three rhesus macaques were enrolled into one of five Cohorts (Fig. 1). Information on the exact timing of infection, ART treatment, and pregnancy for all animals is in Extended Data Table 2. We reasoned that the impacts of SIV co-infection on ZIKV pathogenesis would be most apparent when ZIKV infection occurs early in pregnancy, so we infected animals at approximately GD 30 (range GD 26-38) which corresponds to GD 49 in human pregnancy^28^. Cohort III was SIV naive and not treated with ART to control for the potential impacts of ART on ZIKV adverse outcomes. All SIV+ animals (Cohorts I and IV) reached a chronic set point before beginning ART and achieved undetectable SIV viremia before subsequent ZIKV/mock exposure (Extended Data Fig. 1A). All 14 ZIKV-exposed animals (Cohorts I-III) had detectable ZIKV in plasma by 3 days post inoculation (DPI) and reached peak plasma viremia between 4 DPI and 6 DPI (mean 4.7 DPI) (Extended Data Fig. 1B). Plasma viremia was resolved for all animals by 20 DPI except Animal I who had prolonged detection of ZIKV RNA until 132 DPI; additional data are in Supplementary Table 1. All dams developed robust neutralizing antibody responses to ZIKV by 28 DPI (Supplementary Fig. 1). In total, 11 of 14 (78%) ZIKV+ dams experienced fetal demise in the first trimester (GD 38-54) (Fig. 1).

**Fig 1:**
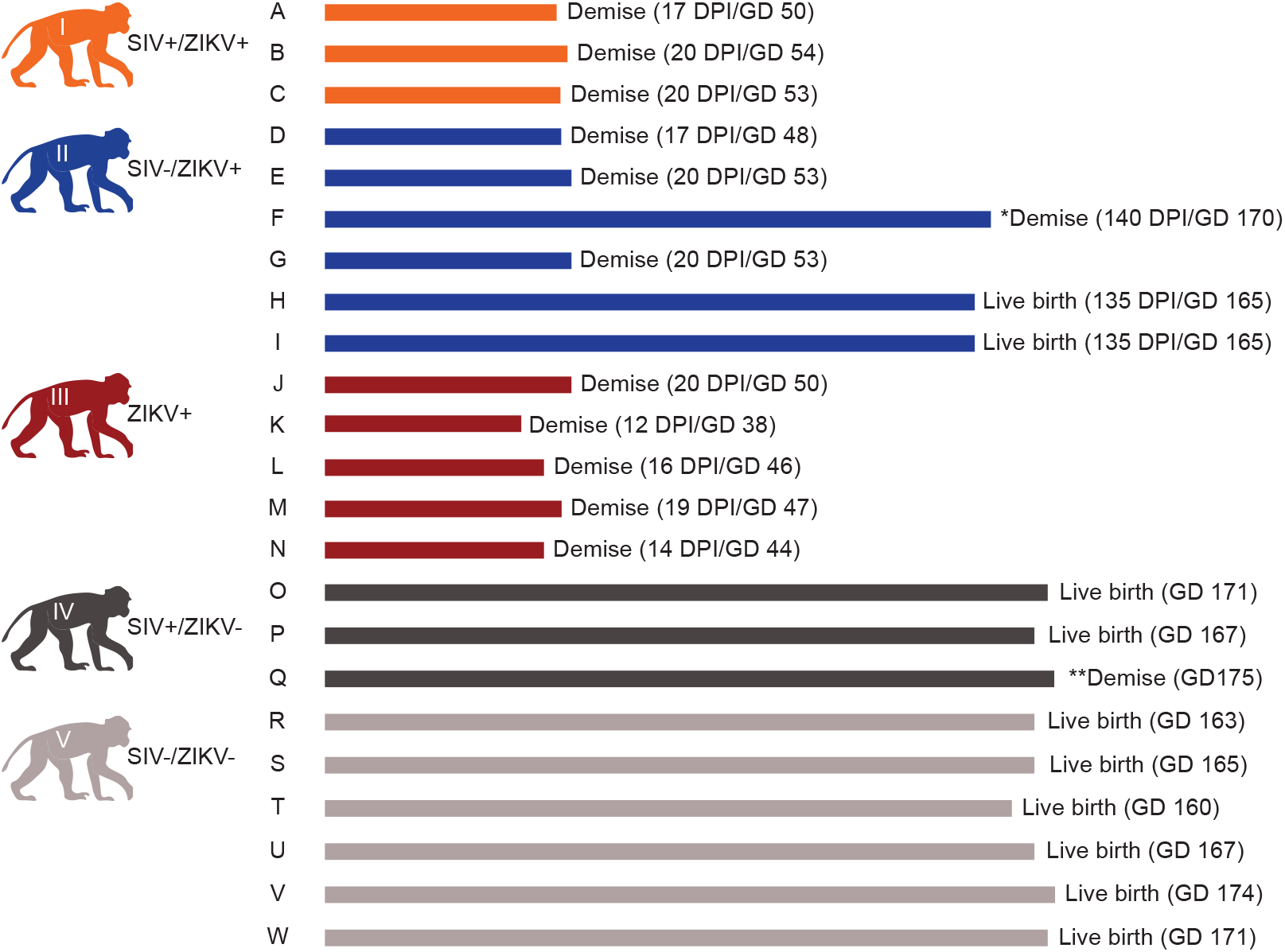
Pregnancy outcomes. *Animal F and fetus died on the day of clinical C-section (GD 170) due to anesthesia complications. **The infant of Animal Q died 5 days after the due date (GD 175).

The timing of pregnancy loss was remarkably consistent: all occurred at 12-20 days post-ZIKV exposure (Fig. 1, Extended Data Table 2). A Cohort II full term infant died from complications (cardiac and respiratory arrest) during clinical Cesarean section (C-section), during which the dam also died (Pregnancy F). More details about this case are in Supplementary Table 2. All animals in the ZIKV naïve cohorts (Cohorts IV and V) maintained viable pregnancies apart from the dam from Pregnancy Q, who experienced a full-term fetal loss around GD 175. This falls within the pregnancy loss rates for non-ZIKV exposed macaques which range from 4-10.9%^14^. In-utero measurements and findings during ultrasounds were recorded and did not identify a consistent phenotype in embryos/fetuses or maternal-fetal interface (MFI) tissue preceding pregnancy loss (Supplementary Fig. 2, Supplementary Tables 3 and 4). Neonatal developmental assessments were performed on surviving infants (Supplementary Figs. 3-9). One of the two infants from Cohort II (Infant H) had lower developmental scores in visual orientation, tracking, and focus tasks in the neonatal period, abnormal retinal function, and a thicker retina compared with the other infants. This suggests that prenatal ZIKV exposure may negatively impact visual pathways in some infants, though larger sample sizes of ZIKV-exposed infants need to be studied to determine if this is a ZIKV-specific impact.

ZIKV was detected in the MFI tissue by reverse transcriptase quantitative polymerase chain reaction (RT-qPCR) from all 11 dams that experienced early pregnancy loss and was present in all three placental layers (decidua, placental parenchyma, chorionic plate) (Fig. 2A). *In situ* hybridization (ISH) also detected ZIKV in the chorionic plate, chorionic villi, and surrounding the chorionic vessels for all 11 cases; however, no RNA was detected in the decidua (Fig. 2B, Supplementary Table 2). The ISH signal in MFI tissue was highly concentrated in the chorionic villi where ZIKV was restricted to the villous stroma and absent from the outer syncytiotrophoblast layer (Fig. 2B, Supplementary Table 2). Histopathologic analysis of placentas from ZIKV-exposed cohorts (Cohorts I-III) identified placental lesions, frequently in the placental parenchyma, in all cases of early pregnancy loss and one case of full-term infant loss (Supplementary Table 2). However, this analysis was limited by the absence of gestational-age matched tissues from ZIKV-naive dams. Nine of 11 dams (Pregnancies A, D, E, G, J, K, L, M, N) across Cohorts I-III had ZIKV detected in a subset of maternal tissues by RT-qPCR (Fig. 2A). ZIKV was also detected in embryonic/fetal tissues and fluids from all cases of early pregnancy loss (Fig. 2C, Fig. 2D, Supplementary Table 2). Pregnancy F had no detectable ZIKV RNA in MFI nor maternal tissues. MFI tissues from Pregnancies H and I were unable to be collected because the dams gave birth naturally.

**Fig 2:**
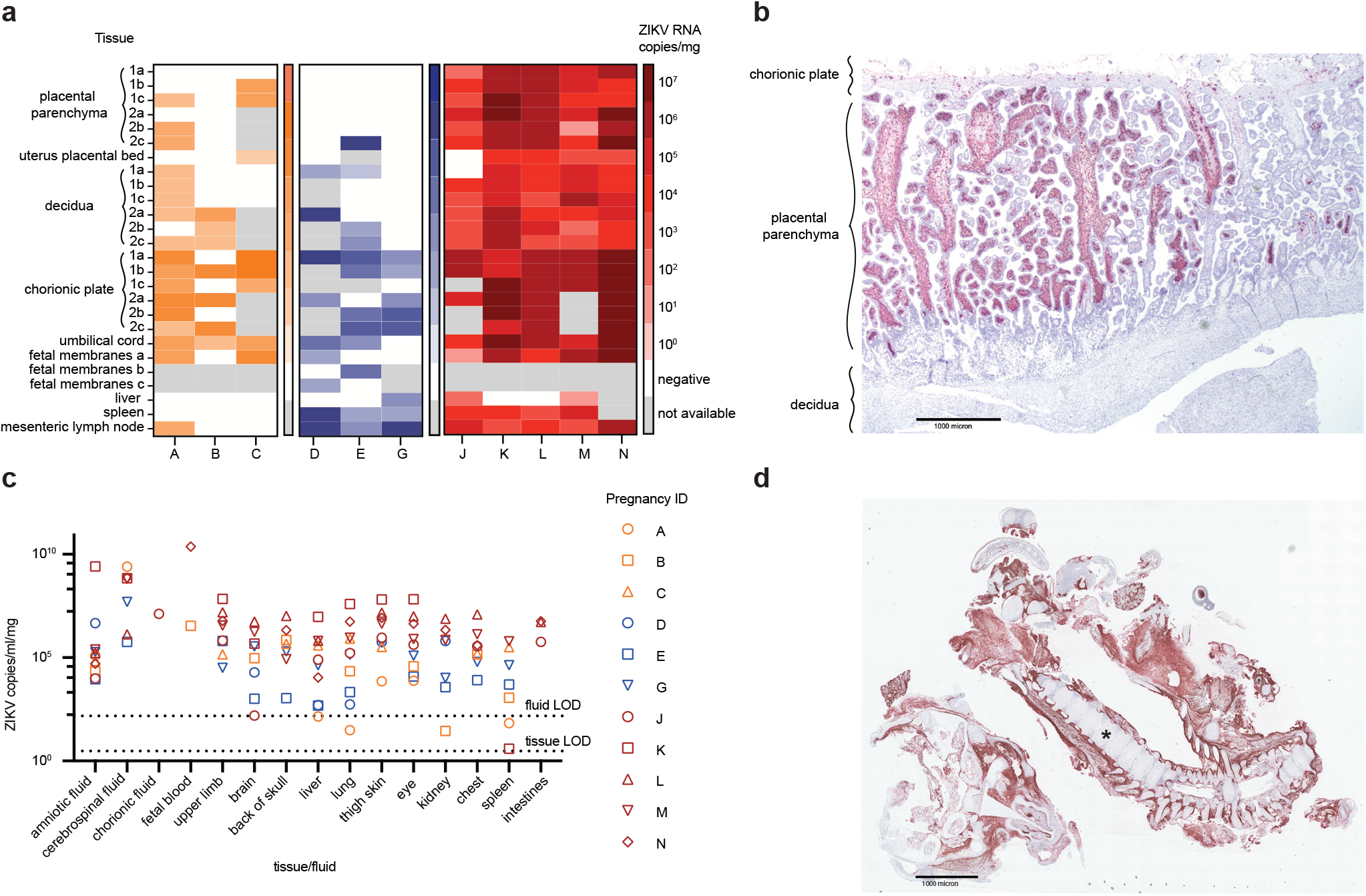
ZIKV was detected in tissues from the maternal-fetal interface (MFI), dam, and fetus/embryo in all cases of early pregnancy loss. Cohort I (SIV+/ZIKV+ +ART) animals are in orange, Cohort II (SIV-/ZIKV+ +ART) animals are in blue, and Cohort III (SIV-/ZIKV+) animals are in red. (a) ZIKV was detected in MFI and maternal tissues from Cohorts I-III by ZIKV-specific RT-qPCR. (b) Representative image of ZIKV RNA (red staining) detected by in-situ hybridization (ISH) in the first placental disc from a case of pregnancy loss (Pregnancy K). Here, there was marked diffuse villous parenchymal staining extending from the basal plate to the chorionic trophoblastic shell with transmural segmental sparing of villi. (c) ZIKV was detected in fetal/embryonic tissues from cases of early pregnancy loss by ZIKV-specific RT-qPCR. LOD denotes the limit of detection. (d) Representative image of ZIKV RNA (red staining) distribution in an embryo from a case of early pregnancy loss (Pregnancy N). Here, ZIKV RNA was detected in the periosteum and musculature of the head and body tissues. The asterisk denotes the vertebral column of the embryo.

With frequent pregnancy loss across all ZIKV+ cohorts (Cohort I-III), we were not powered to detect a difference in pregnancy loss rates in the presence versus absence of SIV coinfection; therefore, we ceased enrollment of animals into our SIV+ Cohorts (I and IV). However, this unexpected finding is arguably more important: we serendipitously developed a model that results in frequent (78%) pregnancy loss in macaques and identified a potentially unappreciated risk for early pregnancy loss in women infected with African-lineage ZIKV.

Previously, macaque infection with the same dose of ZIKV at GD 45 did not result in pregnancy loss^27^. The frequent first-trimester demise that we observe with our GD 30 infection model may be influenced by placental development at the time of infection. ZIKV may be more likely to enter the fetal compartment earlier in gestation due to the remodeling of the spiral arteries^29^. In the first weeks of gestation, fetal extravillous cytotrophoblasts infiltrate maternal decidual spiral arteries, increasing uterine artery blood flow to the placenta^30^. Infection during this critical period may allow more virus access into the fetal compartment and increase the risk of demise. This hypothesis is supported by our earliest ZIKV exposure (Pregnancy K) that occurred at GD 26 and resulted in pregnancy loss earliest (12 DPI). The affected embryo also had the three highest tissue viremia of any pregnancy loss (Fig. 2C).

CZS is a complex phenotype that is likely to be influenced by many factors. In addition to timing of maternal infection, we propose that the difference in pregnancy outcomes noted in different regions of the world may be due to African-lineage ZIKV being more pathogenic than the Asian-lineage viruses that impacted the Americas. Previous studies in mice also suggest a similar conclusion: that African-lineage ZIKV could more easily go unnoticed by public health due to a tendency to cause fetal loss rather than birth defects^24,25^. Translating our pregnancy loss rate of 78% in macaques to humans, it is possible that CZS and microcephaly have gone unreported in Africa because ZIKV infections frequently result in miscarriage, possibly before the pregnancy is recognized. Our study in macaques is impactful because the similarities to humans in placental development and immunology make this model particularly translational.

Further studies focusing on mechanisms of the fetal demise phenotype are needed to fully understand the adverse pregnancy outcomes we observed and develop effective countermeasures. There is also the possibility that the ZIKV isolate used in this study is not broadly representative of viruses currently circulating throughout sub-Saharan Africa, however, there are more contemporary, geographically representative ZIKV isolates available from reference centers to perform comparative analyses^24^. Additionally, a sustained surveillance effort in African populations will be important to understand if African ZIKV is a looming threat for global health.

There are currently no FDA-approved countermeasures for ZIKV infection (https://www.fda.gov/emergency-preparedness-and-response/mcm-issues/zika-virus-response-updates-fda), in part due to waning ZIKV outbreaks and the absence of a translational pregnancy model that results in consistent outcomes to assess medical countermeasures. Consistent outcomes are needed to make robust comparisons in macaque studies that are inherently limited by small sample sizes. The first-trimester African-lineage ZIKV exposure model described here provides new opportunities for testing therapeutics.

## Online Methods

### Care and use of macaques

The macaques used in this study were cared for by the staff at the Wisconsin National Primate Research Center (WNPRC) in accordance with recommendations of the Weatherall report and the principles described in the National Research Council’s Guide for the Care and Use of Laboratory Animals^31^. The University of Wisconsin - Madison, College of Letters and Science and Vice Chancellor for Research and Graduate Education Centers Institutional Animal Care and Use Committee approved the nonhuman primate research under protocol number G006139. The University of Wisconsin - Madison Institutional Biosafety Committee approved this work under protocol number B00000117. Animals were housed in enclosures with the required floor space and fed using a nutritional plan based on recommendations published by the National Research Council. Animals were fed an extruded dry diet with adequate carbohydrates, energy, fat, fiber, mineral, protein, and vitamin content. Diets were supplemented with fruits, vegetables, and other edible objects (e.g., nuts, cereals, seed mixtures, yogurt, peanut butter, popcorn, marshmallows, etc.) to provide variety and to inspire species-specific behaviors such as foraging. To promote psychological well-being, animals were provided with food enrichment, structural enrichment, and/or manipulanda. Environmental enrichment objects were selected to minimize the chances of pathogen transmission from one animal to another and from animals to care staff. While on the study, all animals were evaluated by trained animal care staff at least twice daily for signs of pain, distress, and illness by observing appetite, stool quality, activity level, and physical condition. Animals exhibiting abnormal presentation for any of these clinical parameters were provided appropriate care by attending veterinarians. Before all minor/brief experimental procedures, macaques were sedated using ketamine anesthesia and regularly monitored until fully recovered.

### Study design

Twenty-three female rhesus macaques (*Macaca mulatta)* were divided into five cohorts denoted Cohort I through Cohort V (Table 2). Cohorts I (SIV+/ZIKV+ +ART), II (SIV-/ZIKV+ +ART), IV (SIV+/ZIKV-+ART), and V (SIV-/ZIKV-+ART) were exposed to 300 TCID_50_ SIV-mac239 (SIV+) or mock (SIV-) with 1xPBS intrarectally (IR). The dam from Pregnancy O (Cohort IV) was not successfully infected with SIV. Thus, this animal was re-exposed to 500TCID_50_ SIV-mac239 intravenously 21 days after IR exposure. These four cohorts were treated once daily with injectable combination ART (+ART) consisting of tenofovir disoproxil fumarate (TDF), emtricitabine (FTC), and dolutegravir sodium (DTG) (see **Antiretroviral therapy**). Once treated SIV+ animals controlled viremia under the limit of quantification of our in-house SIV RT-qPCR assay (<200 copies/ml plasma, see **Viral RNA quantification by RT-qPCR**), animals had their combination ART switched to an injectable combination of ART of TDF and FTC with two oral doses of Raltegravir (RAL, 100mg/dose) for 30 days before and throughout breeding (see **Antiretroviral therapy**). All five cohorts underwent timed breeding until pregnancy was confirmed by ultrasound. Animals maintained combination ART (TDF/FTC/RAL) throughout pregnancy. At approximately GD 30, animals in Cohorts I, II, and III, were subcutaneously (SC) exposed to 1×10^4^ plaque forming units (PFU)/ml of African-lineage ZIKV (ZIKV+), while Cohorts IV and V were SC exposed to 1xPBS (ZIKV-). Animals were enrolled in Cohort III (ZIKV+ -ART) to confirm that adverse pregnancy outcomes in Cohort I (SIV+/ZIKV+ +ART) and Cohort II (SIV-/ ZIKV+ +ART) were the result of ZIKV exposure and not additive impact from ART treatment. All pregnancies were monitored throughout the study with weekly ultrasounds and plasma vRNA load quantification of ZIKV and SIV where appropriate. Pregnancies were allowed to go to term and natural delivery; however, a C-section was performed in the event of an overdue pregnancy (GD 175) (Pregnancies F, O, Q and W) or demise (no detection of fetal/embryonic heartbeat). In cases of demise, the C-section was followed by a fetal or embryonic necropsy, maternal biopsies, and MFI tissue collection.

### Antiretroviral therapy

Animals in Cohorts I, II, IV, and V were treated daily with an injectable combination ART of sterile-filtered TDF (final concentration 5.1mg/ml), FTC (final concentration 50mg/ml), and DTG (final concentration 2.5mg/ml) in the commercially-available solubility vehicle Kleptose (Roquette, Gurnee, IL). ART drugs were sourced from Hangzhou APIChem Technology Co., Ltd. (Hangzhou, Zhejiang, China) and were confirmed by mass-spectrophotometry at the University of Wisconsin-Madison Genetics and Biotechnology Center. This combination of ART drugs, which includes two nucleoside reverse transcriptase inhibitors (TDF and FTC) and an integrase inhibitor (DTG), has been previously shown to control SIV infection in macaques when provided as a combination injectable at a dose of 1ml/kg^32,33^. Beginning 30 days before breeding and then continuing throughout pregnancy, animals were given oral doses of raltegravir (RAL, 100mg/dose) twice daily alongside a modified daily injection containing TDF and FTC (5.1mg/ml and 50mg/ml, respectively). The integrase inhibitor RAL was used in place of DTG at this study stage due to the potential association of DTG with neural tube defects in human infants when used during pregnancy^34^. Following birth, animals transitioned from oral RAL/double-combination injectable back to triple-combination injectable for continued maintenance of SIV infections. Mock-SIV animals stopped ART after the births of their infants.

### Ultrasonography and fetal monitoring

Ultrasounds and fetal doppler were conducted weekly (Cohorts I, II, IV, and V) to observe the growth and viability of the fetus and to obtain measurements including fetal femur length (FL), biparietal diameter (BPD), head circumference (HC), and heart rate as previously described ^12,35^. Mean growth measurements were plotted against mean measurements and standard deviations from specific gestational days collected from rhesus macaques^36^. Comparison of experimental growth parameters with the established growth curves allowed extrapolation of actual gestational age versus predicted gestational age^12^. The standard growth curve was extrapolated to contextualize measurements collected before GD 50. For Cohort III, fetal dopplers were performed more frequently (daily from 10-21 days post-ZIKV infection) to confirm viability.

### ZIKV Infection

Zika virus strain Zika virus/A.africanus-tc/Senegal/1984/DAKAR 41524 (ZIKV-DAK; GenBank: KX601166, SRR7879856) was originally isolated from *Aedes luteocephalus* mosquitoes in Senegal in 1984. One round of amplification on *Aedes pseudocutellaris* cells, followed by amplification on C6/36 cells, followed by two rounds of amplification on Vero cells, was performed by BEI Resources (Manassas, VA) to create the stock ^25^. Once obtained, an additional expansion was performed on C6/36 cells. Stocks used to infect the animals were prepared from three different passages and sequencing showed the stock viruses to be identical at the consensus levels. No minor variants were present at >10% in any of the stocks. For virus challenges, ZIKV-DAK stock was diluted to 1×10^4^ PFU in 1ml of 1x phosphate buffered saline (1x PBS) and delivered to each dam SC over the cranial dorsum via a 1ml luer lock syringe.

### SIV Infection

Simian immunodeficiency virus (SIV-mac239, Genbank: <u>M33262</u>) stock was produced from two plasmids acquired from the AIDS Reagent Resource. Plasmids were ligated and transfected in E6 Vero cells. Cell-derived supernatant was then used to infect cultured macaque CD8+ peripheral blood mononuclear cells (PBMC), which were then monitored for virus production. The supernatant was harvested at peak virus production. SIV-mac239 was initially used to intra-rectally (IR) expose all animals in Cohorts I and IV (A, B, C, P/Q, and O) at a dose of 300 TCID_50_. Following initial IR exposure, Animal O was found to be uninfected and was subsequently re-exposed intravenously with 500 TCID_50_ with the same virus stock 21 days later. All virus stock dilutions were made in sterile-filtered 1x PBS and administered in a 1ml syringe.

### Blood and body fluids monitoring

Blood samples were collected for isolation of plasma and PBMC from dams prior to SIV infection on days -1 and 0, post-SIV infection on days 7, 13, 14, 16, weekly through 4 weeks post-infection, and twice weekly until ZIKV infection. Blood samples, urine, and saliva were collected on days -1, 0, 3-7, 10, 14 post-ZIKV challenge, and then twice weekly until 28 DPI or until ZIKV was undetectable in blood plasma by RT-qPCR. Samples were then collected weekly until birth.

### Viral RNA isolation from plasma, urine, and saliva

Plasma and PBMC were isolated from EDTA-treated whole blood by layering blood on top of ficoll in a 1:1 ratio and performing centrifugation at 1860x rcf for 30 minutes with no brake. Plasma and PBMC were extracted and transferred into separate sterile tubes. R10 medium warmed at 37 degrees Celsius was added to PBMC before a second centrifugation of both tubes at 670 x rcf for 8 minutes. Before treatment, media was removed from PBMC with 1x Ammonium-Chloride-Potassium (ACK) lysing buffer for 5 minutes to remove red blood cells. An equal amount of R10 medium was added to quench the reaction before another centrifugation at 670 x rcf for 8 minutes. Supernatant was removed before freezing down of cells in CryoStor CS10 medium (BioLife Solutions, Inc., Bothell, WA) for long-term storage in liquid nitrogen freezers. Serum was obtained from clot activator tubes by centrifugation at 670 x rcf for 8 minutes or from serum separation tubes (SST) at 1400 x rcf for 15 minutes. Urine was passively collected from the bottom of animals’ housing, centrifuged for 5 minutes at 500 x rcf to pellet debris, and 270ul was added into 30ul dimethyl sulfoxide (DMSO) followed by slow freezing. Saliva swabs were obtained and put into 500ul viral transport media (VTM) consisting of tissue culture medium 199 supplemented with 0.5% FBS and 1% antibiotic/antimycotic. Tubes with swabs were vortexed and centrifuged at 500 x rcf for 5 minutes. Viral RNA (vRNA) was extracted from 300uL plasma, 300uL saliva+VTM, or 300ul urine+DMSO using the Maxwell RSC Viral Total Nucleic Acid Purification Kit on the Maxwell 48 RSC instrument (Promega, Madison, WI).

### Maternal, fetal, and maternal-fetal interface tissue (MFI) collection from first-trimester pregnancy losses

Following early pregnancy loss, fetal, maternal, and MFI tissues were harvested by board certified veterinary pathologists at the WNPRC. Recovered MFI tissues for pathological evaluation included three sections from each placental disc, amniotic/chorionic membrane from each placental disc, decidua from each placental disc, and one section from the decidua parietalis (fetal membranes), umbilical cord, and uterus/placental bed. One section of each of the following maternal or fetal tissues was also collected: maternal liver, maternal spleen, mesenteric lymph node (LN), fetal liver, fetal intestine, fetal lung, fetal kidney, fetal brain, fetal skin/muscle from thigh, fetal eye, fetal spleen, fetal upper limb, fetal chest, and fetal skull with brain. Two samples from each tissue section were collected and stored in either 750ul VTM or 1mL RNAlater for vRNA assessment and future analysis. Tissues in VTM were frozen immediately after collection and stored at -80°C. Tissues in RNAlater were refrigerated for 24 hours at 4°C, after which RNAlater was aspirated off, and the tissues were stored at -80°C prior to vRNA isolation.

### SIV RNA quantification by RT-qPCR

Viral RNA was quantified using an RT-qPCR assay based on the one developed by Cline et al.^37^. RNA was reverse transcribed and amplified using the TaqMan Fast Virus 1-Step Master Mix RT-qPCR kit (Invitrogen) on the LightCycler 480 instrument (Roche, Indianapolis, IN), and quantified by interpolation onto a standard curve made up of serial ten-fold dilutions of in vitro transcribed RNA. RNA for this standard curve was transcribed from the p239gag_Lifson plasmid kindly provided by Dr. Jeffrey Lifson, NCI/Leidos. The final reaction mixtures contained 150 ng random primers (Promega, Madison, WI), 600 nM each primer, and 100 nM probe. Primer and probe sequences are as follows: forward primer: 5’-GTCTGCGTCATPTGGTGCATTC-3, reverse primer:5′-CACTAGKTGTCTCTGCACTATPTGTTTTG-3′ and probe:5′-6-carboxyfluorescein-CTTCPTCAGTKTGTTTCACTTTCTCTTCTGCG-BHQ1-3’. The reactions cycled with the following conditions: 50°C for 5 minutes, 95°C for 20 seconds followed by 50 cycles of 95°C for 15 seconds and 62°C for 1 min. The limit of detection of this assay is 200 copies/ml.

### ZIKV RNA isolation from tissue samples

Isolation of RNA from tissue samples was performed using a modification of the method described by Hansen, et al.^38^. Up to 200mg of tissue was disrupted in TRIzol Reagent (Thermo Fisher Scientific, Waltham, MA) with stainless steel beads (2×5 mm) using a TissueLyser (Qiagen, Germantown, MD) for three minutes at 25 r/s twice. Following homogenization, samples in TRIzol were separated using bromo-chloro-propane (Sigma, St. Louis, MO). The aqueous phase was collected into a new tube and glycogen was added as a carrier. The samples were washed in isopropanol and ethanol-precipitated overnight at -20°C. RNA was then fully re-suspended in 5 mM Tris pH 8.0.

### ZIKV RNA quantification by RT-qPCR

Viral RNA was quantified using a highly sensitive RT-qPCR assay based on the one developed by Lanciotti et al.^39^, though the primers were modified to accommodate both Asian and African lineage ZIKV lineages. RNA was reverse transcribed and amplified using the TaqMan Fast Virus 1-Step Master Mix RT-qPCR kit (Invitrogen) on a LightCycler 480 or LC96 instrument (Roche, Indianapolis, IN), and quantified by interpolation onto a standard curve made up of serial tenfold dilutions of in-vitro transcribed RNA. RNA for this standard curve was transcribed from a plasmid containing an 800 base pair region of the ZIKV genome targeted by the RT-qPCR assay. The final reaction mixtures contained 150 ng random primers (Promega, Madison, WI), 600 nM each primer and 100 nM probe. Primer and probe sequences are as follows:

forward primer: 5’-CGYTGCCCAACACAAGG-3’

reverse primer: 5′-CCACYAAYGTTCTTTTGCABACAT-3′

and probe: 5′-6-carboxyfluorescein-AGCCTACCTTGAYAAGCARTCAGACACYCAA-BHQ1-3’. The reactions cycled with the following conditions: 50°C for 5 minutes, 95°C for 20 seconds followed by 50 cycles of 95°C for 15 seconds, and 60°C for 1 min. The limit of detection of this assay in body fluids is 150 copies/ml and 3 copies/mg in tissues.

### IgM ELISA

An IgM ELISA was performed on serum samples collected on days 0, 7, 13, and 21 following ZIKV infection. Samples were run in triplicate using the AbCam anti-Zika virus IgM micro-capture ELISA kit protocol according to the manufacturer’s instructions (cat# ab213327, Abcam Inc., Cambridge, UK). Briefly, samples were thawed to room temperature, added to an anti-human IgM-coated microplate tray (μ capture), and incubated. Zika virus conjugate+HRP was added, followed by TMB substrate solution (3, 3’, 5, 5’-tetramethylbenzidine < 0.1%), and stop solution (sulphuric acid, 0.2 mol/L). The plate absorbance was read at dual wavelengths of 450nm and 600nm within 30 minutes of adding the stop solution, and the IgM concentration was measured in the calculated Abcam units (AU) relative to the kit cut-off control. To calculate the AU, the 600nm well data were first subtracted from the 450nm well data. Because multiple samples were run for each animal at each DPI, the average of the numbers was calculated, multiplied by ten, and divided by the absorbance of the cut-off control to get a single AU value per sample. Samples were considered positive if they were above 10 AU and negative if they were below 10 AU.

### Plaque Reduction Neutralization Test

Titers of ZIKV neutralizing antibodies (nAb) were determined for days 0, 21, or 28 post-ZIKV infection using PRNT on Vero cells (ATCC #CCL-81) with a cutoff value of 90% (PRNT^90^)^40^. Briefly, ZIKV-DAK was mixed with serial 2-fold dilutions of serum for 1 hour at 37°C prior to being added to Vero cells. Neutralization curves were generated in GraphPad Prism (San Diego, CA) and the resulting data were analyzed by nonlinear regression to estimate the log_10_ reciprocal serum dilution required to inhibit 90% infection of Vero cell culture^40,41^.

### *In situ* hybridization (ISH)

ISH probes against the ZIKV genome were commercially purchased (cat# 468361, Advanced Cell Diagnostics, Newark, CA). ISH was performed using the RNAscope® Red 2.5 kit (cat# 322350, Advanced Cell Diagnostics, Newark, CA) according to the manufacturer’s protocol. After deparaffinization with xylene, a series of ethanol washes, and peroxidase blocking, sections were heated with the antigen retrieval buffer and then digested by proteinase. Sections were then exposed to the ISH target probe and incubated at 40°C in a hybridization oven for two-hours. After rinsing, the ISH signal was amplified using the provided pre-amplifier followed by the amplifier-containing labeled probe binding sites, and developed with a Fast Red chromogenic substrate for 10 minutes at room temperature. Sections were then stained with hematoxylin, air-dried, and mounted.

### Statistical analyses

We defined peak plasma viremia as the highest ZIKV plasma viremia detected for each dam in Cohorts I-III. Plasma viremia duration was defined for these animals as the last time point a dam had ZIKV detected in plasma by RT-qPCR above the limit of quantification of the assay. Overall plasma ZIKV RNA loads were calculated for all ZIKV-infected dams (Cohorts I-III) using the trapezoidal method to calculate AUC in R Studio v. 1.4.1717. AUC values were then compared between Cohorts I-III using a Kruskall-Wallis rank sum test. Peak plasma ZIKV RNA loads, as well as the duration of positive ZIKV RNA detection, were also compared between Cohorts I-III using a Kruskall-Wallis rank sum test (duration) and one-way ANOVA (peak plasma viremia) using R Studio v. 1.4.1717. The time to peak plasma ZIKV RNA load was also compared between dams in Cohorts I-III. Time to peak was analyzed using a one-way ANOVA to compare between Cohorts. For all analyses of plasma ZIKV RNA loads, p<0.05 was used to define statistical significance. In-utero growth trajectories of abdominal circumference (AC), biparietal diameter (BPD), femur length (FL), and head circumference (HC) were quantified by fitting a linear mixed-effects regression model with animal-specific random effects and an autoregressive correlation structure over time, in this case, weeks post-infection (WPI) using SAS version 9.4 (SAS Institute, Cary NC). Since the growth trajectories were non-linear, a log transformation for each outcome was used. Growth trajectories were compared between Cohorts I, IV and V by comparing the corresponding slopes (Supplementary Table 3) and graphs were generated using R software v. 4.1.0 (R Foundation for Statistical Computing) (Supplementary Fig. 2). No statistical analyses of infant development, vision, and hearing tests were performed due to small group sizes.

### Infant developmental tests

The Schneider Neonatal Assessment for Primates (SNAP) was used to assess the neurodevelopmental areas of interest (Orientation, Motor maturity and activity, Sensory responsiveness, and State control). This neonatal test is well validated and has previously been used to define neonatal development of prenatally ZIKV-exposed infants^16^. The Catwalk XT version 10.6 was modified for infant rhesus macaques and used to assess gait development, as described previously^26,42^. The SNAP was administered at 7, 14, 21, and 28 (+/-2) days of life, and the Catwalk was administered on 14, 21, and 28 days of life.

### Infant vision and hearing tests

Infants were anesthetized for eye exams performed by a human ophthalmologist with retinal fellowship training (M. Nork). Slit-lamp biomicroscopy and indirect ophthalmoscopy were performed after pupillary dilation. To evaluate visual function, standard visual electrodiagnostic procedures including a full-field electroretinogram (ERG) and the cortical-derived visual evoked potential (VEP) were performed as previously described^13^. To define retinal layer structure, spectral-domain optical coherence tomography (OCT) was performed as previously described^13^. Auditory brainstem response (ABR) testing was completed, which measures brainstem evoked potentials generated by a brief click, 500 Hz stimulus, or 1000 Hz stimulus, as previously described. The presence or absence of a Wave IV response was recorded for each decibel level and stimulus^13^. The presence or absence of a Wave IV response was recorded for each decibel level and stimulus.

## Data Availability

All relevant data are within the manuscript and the Supplementary Information files. Primary data that support this study are also available at GitHub https://github.com/lraasch/Frequent-first-trimester-pregnancy-loss-in-rhesus-macaques-infected-with-African-lineage-Zika-virus. ISH images are available at https://go.https://go.wisc.edu/23d838wisc.edu/23d838.

## Acknowledgments

This work was supported by NIH Grants R01AI138647, P01AI132132, K08AAI3398, R01AAH9849,and P51OD011106. This work was supported in part by an Unrestricted Grant from Research to Prevent Blindness, Inc. to the UW-Madison Department of Ophthalmology and Visual Sciences and in part by the Core Grant for Vision Research from the NIH to UW-Madison (P30 EY016665). Authors thank the McPherson Eye Research Institute at UW-Madison. Authors also thank Max C. Ertl, Mitchell D. Ramuta, and Ryan V. Moriarty for their thoughtful reading and discussion.

## Notes

https://github.com/lraasch/Frequent-first-trimester-pregnancy-loss-in-rhesus-macaques-infected-with-African-lineage-Zika-virus

https://go.wisc.edu/23d838

### Competing Interest Statement

The authors have declared no competing interest.

