## Supplemental_tables_and_figures for "Frequent first-trimester pregnancy loss in rhesus macaques infected with African-lineage Zika virus"

**Supplemental Table 1: Demise rate of Asian-lineage pregnancy studies in macaques.** Across multiple studies the demise rate was <10% when animals were exposed to an Asian-lineage strain of Zika virus, PRVABC59.

| Animal | Infection GD | Pregnancy outcome |
|---|---|---|
| Dam[16] | 45 | Live Birth |
| Dam[16] | 45 | Live Birth |
| Dam[16] | 45 | Live Birth |
| Dam[16] | 45 | Live Birth |
| Dam[16] | 45 | Live Birth |
| Dam[15] | 30 | Live Birth |
| Dam[15] | 25 | Live Birth |
| Dam[15] | 26 | Live Birth |
| Dam[15] | 30 | Live Birth |
| Dam[15] | 30 | Live Birth |
| Dam[15] | 33 | Live Birth |
| Dam[17] | 45 | Live Birth |
| Dam[17] | 45 | Live Birth |
| Dam[17] | 45 | Live Birth |
| Dam[17] | 45 | Live Birth |
| Dam[12] | 46 | Demise |
| Dam[13] | 45 | Live Birth |
| Dam[13] | 45 | Live Birth |
| Dam[13] | 45 | Live Birth |
| Dam[13] | 45 | Live Birth |
| Dam[13] | 45 | Demise |

**Supplemental Table 2: Correspondent demographic information by Cohort and Pregnancy ID throughout the study period.** Age and weight presented correspond to the measures taken at the ZIKV challenge date. GD denotes gestational day and DPI denotes days post infection. Animals with pregnancies that made it to near-term delivered naturally (N) or by Cesarean section (C). In all cases of pregnancy loss the fetus/embryo was extracted by Cesarean section (C).

| | | | | | **ZIKV challenge** | | | | |
|---|---|---|---|---|---|---|---|---|---|
| **Cohort** | **Pregnancy ID** | **Age (years)** | **Weight (kg)** | **Date of SIV challenge** | **Date** | **GD at challenge** | **GD at demise** | **DPI at demise/birth** | **Full-term pregnancy GD** |
| I. SIV+/ZIKV+ +ART | A | 10 | 8.84 | 2020-02-04 | 2020-10-09 | 33 | 53 (C) | 20 | |
| | B | 6 | 6.78 | 2020-02-04 | 2020-09-11 | 34 | 54 (C) | 20 | |
| | C | 11 | 9.76 | 2020-02-04 | 2020-09-11 | 33 | 50 (C) | 17 | |
| II. SIV-/ZIKV+ +ART | D | 7 | 6.54 | 2019-08-21 | 2020-02-07 | 31 | 48 (C) | 17 | |
| | E | 15 | 6.08 | 2019-08-21 | 2020-04-24 | 33 | 53 (C) | 20 | |
| | F | 7 | 8.85 | 2020-01-15 | 2020-06-19 | 35 | 175 (C) | 140 | |
| | G | 11 | 6.91 | 2020-01-15 | 2020-07-24 | 33 | 53 (C) | 20 | |
| | H | 17 | 7.09 | 2020-11-17 | 2021-06-11 | 30 | | 135 | 165N) |
| | I | 7 | 6.30 | 2020-11-17 | 2021-06-04 | 29 | | 135 | 164 (N) |
| III. ZIKV+ | J | 6 | 6.87 | | 2021-11-12 | 30 | 50 (C) | 20 | |
| | K | 5 | 5.53 | | 2022-01-28 | 26 | 38 (C) | 12 | |
| | L | 5 | 6.26 | | 2022-02-04 | 30 | 46 (C) | 16 | |
| | M | 6 | 7.72 | | 2022-04-01 | 27 | 46 (C) | 19 | |
| | N | 15 | 5.71 | | 2022-05-27 | 32 | 46 (C) | 14 | |
| IV. SIV+/ZIKV- +ART | O | 7 | 10.42 | 2020-02-04 | 2021-12-17 | 28 | | 143 | 171 (C) |
| | P | 8 | 6.30 | 2020-02-04 | 2020-10-01 | 38 | | 136 | 174 (N) |
| | Q | 10 | 7.07 | 2020-02-04 | 2022-05-27 | 29 | 175 (C) | 141 | |
| V. SIV-/ZIKV- +ART | R | 15 | 8.29 | 2019-08-21 | 2020-01-24 | 30 | | 135 | 165 (N) |
| | S | 16 | 9.36 | 2019-08-21 | 2020-03-06 | 29 | | 137 | 166 (N) |
| | T | 13 | 7.10 | 2020-01-15 | 2020-10-09 | 31 | | 131 | 162 (N) |
| | U | 15 | 6.42 | 2020-11-17 | 2021-05-07 | 31 | | 135 | 166 (N) |
| | V | 15 | 11.08 | 2020-11-17 | 2021-07-23 | 30 | | 141 | 171 (N) |
| | W | 11 | 6.74 | 2020-11-17 | 2021-06-04 | 27 | | 147 | 174 (C) |

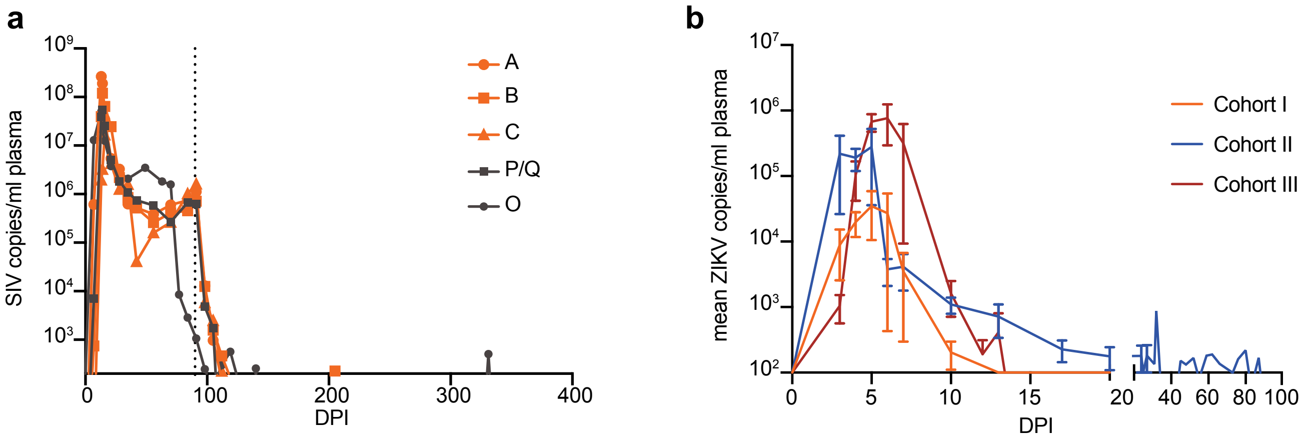


**Supplemental Fig. 1: Maternal plasma viremia.** (a) SIV viremia in macaque plasma before and after ART regimen. Copies of viral RNA were determined by SIV-specific RT-qPCR. Cohort I (SIV+/ZIKV+ +ART) pregnancies are in orange, Cohort IV (SIV+/ZIKV- +ART) pregnancies are in dark gray. The dotted line at 90 DPI indicates when animals started the 1x daily injectable combination ART regimen (TDF/FTC/DTG). Pregnancies P and Q are from the same dam and thus have the same SIV plot lines depicted here. (b) ZIKV viremia in macaque plasma. Copies of viral RNA were detected by ZIKV-specific RT-qPCR. Mean ZIKV plasma viremia are displayed in orange for Cohort I, blue for Cohort II (SIV-/ZIKV+ +ART), and red for Cohort III (SIV-/ZIKV+). Bars represent the standard error of the mean (SEM).

**Supplemental Table 3: Statistical analysis of ZIKV kinetics in macaque plasma.** Viremia was measured by ZIKV-specific RT-qPCR. Area under the curve for plasma viremia, duration of plasma viremia, magnitude of peak plasma viremia, and time from ZIKV exposure to peak plasma viremia was compared between Cohorts I-III. None of the analyses revealed significant differences between groups.

| | Test | Results |
|---|---|---|
| Area under the curve | Kruskal-Wallis rank sum | $X_2 = 5.88$, df = 2, p = 0.053 |
| Duration | Kruskal-Wallis | $X_2 = 3.66$, df = 2, p = 0.16 |
| Magnitude of peak viremia | One-way ANOVA | $F(2, 11) = 3.06$, p = 0.09 |
| Time to peak viral load | One-way ANOVA | $F(2,11) = 2.36$, p = 0.14 |

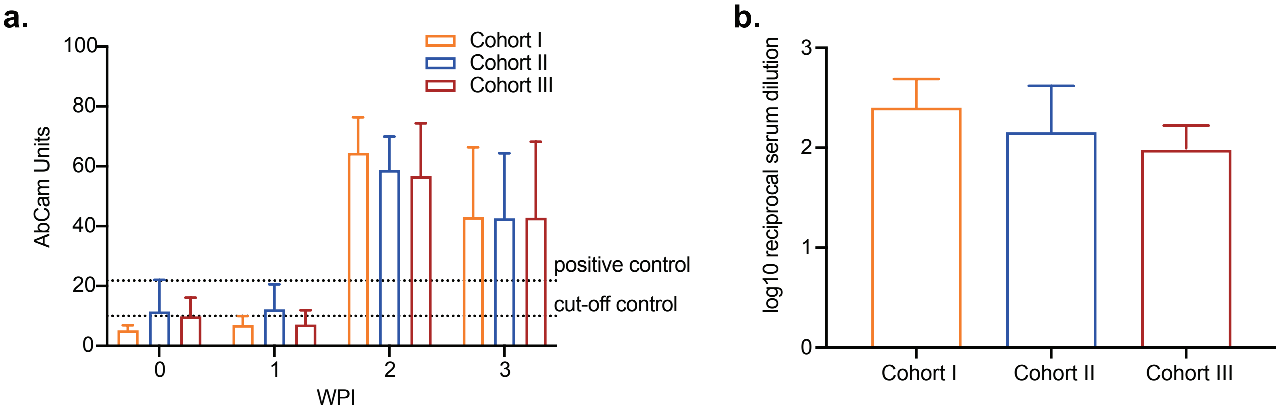


**Supplemental Fig. 2: All ZIKV-exposed animals developed robust IgM and neutralizing antibody responses** (a) ELISA determined Anti-ZIKV IgM levels at 0, 1, 2, and 3 weeks post infection (WPI). (b) Neutralizing antibody levels were measured at 4 WPI by PRNT 90.

**Supplemental Table 4. Histological diagnosis and localization of ZIKV RNA in placental and embryonic/fetal tissue from cases of early pregnancy loss.** Lesions are graded on a scale of 0-4: 0=no inflammation/pathology/necrosis; 1=minimal; 2=mild; 3=moderate; 4=severe. Localization of ZIKV RNA was determined by in-situ hybridization (ISH).

| Cohort | Pregnancy ID | GD | Pathology and ISH remarks |
|---|---|---|---|
| I | A | 53 | Uterus: multifocal neutrophilic endometritis (1).<br>Decidua basalis: multifocal acute deciduitis with subacute hemorrhage and hemosiderosis (2).<br>Decidua parietalis: (0).<br>Placental parenchyma: multifocal chronic villitis, acute intervillositis, multifocal mineralization, diffuse chorionic hemorrhage, and multifocal basal plate thrombosis (2).<br>Umbilical cord: (0).<br>ISH - Placenta, center cuts (Disc 1 and 2): scattered positive staining within the villous parenchyma, the chorionic plate, and multiple chorionic vessels for both placental discs (3).<br>Fetus: diffuse autolysis (4) and multifocal interstitial pulmonary hemorrhage with rare single neutrophils (3).<br>ISH - Fetus: minimal positive staining of the skin and meninges surrounding the neuropil (cerebrum). There is minimal multifocal scattered single cell positive staining of skeletal muscle and subcutaneous tissue of the hind limbs. |

| | | | | |
|---|---|---|---|---|
| 87 | I | B | 54 | Uterus: Nc.<br>Decidua basalis: (0).<br>Decidua parietalis. (0).<br>Placental parenchyma: mild multifocal chorionic vascular thrombosis, mild multifocal villous stromal karyorrhectic debris (VSVK), intervillous syncytial knots, multifocal villus ischemia, focal hemorrhage, and minimal basal plate infarction (2)..<br>ISH Placenta, center cut Disc 1: intense central staining of most placental villi, minimal to mild scattered diffuse single cell staining within the chorionic plate, minimal to mild staining within the wharton's jelly of the umbilical cord.<br>ISH cotyledon center cut Disc 2: There is minimal scattered positive staining within the chorionic plate surrounding chorionic vessels, occasionally more intense staining within the stroma of scattered single placental villi.<br>Umbilical cord: (0).<br>Fetus: diffuse autolysis (3).<br>ISH - Fetus: intense positive staining within the outer muscular layer and serosa of multiple sections of intestine, mild scattered single cells staining within the intestinal mesentery, scattered single cell staining within within the renal interstitium with intense positive staining of the periosteum of long bones and adjacent skeletal muscle as well as scattered single cell staining within the dura mater, nasal turbinates, and choroid plexus. |
| 88<br>89 | I | C | 50 | Uterus: (0).<br>Decidua basalis: (0).<br>Decidua parietalis: (0).<br>Placental parenchyma: mild multifocal chorionic vascular thrombosis, mild multifocal villous stromal karyorrhectic debris (VSVK), intervillous syncytial knots, multifocal villus ischemia, focal hemorrhage, and minimal basal plate infarction (2).<br>IISH Placenta, center cut Disc 1: intense central staining of most placental villi, minimal to mild scattered diffuse single cell staining within the chorionic plate, minimal to mild staining within the wharton's jelly of the umbilical cord.<br>Umbilical cord: (0).<br>Fetus: diffuse autolysis (3).<br>ISH - Fetus: intense positive staining within the outer muscular layer and serosa of multiple sections of intestine, mild scattered single cells staining within the intestinal mesentery, scattered single cell staining within within the renal interstitium with intense positive staining of the periosteum of long bones and adjacent skeletal muscle as well as scattered single cell staining within the dura mater, nasal turbinates, and choroid plexus. |

| | | | | |
|---|---|---|---|---|
| 90 | II | D | 48 | Uterus: Moderate multifocal lymphoplasmacytic and mildly neutrophilic deciduitis (3).<br>Decidua basalis: Multifocal, vasculocentric, necrotizing deciduitis (3) with neutrophilic and lymphoplasmacytic infiltration (2).<br>Decidua parietalis: multifocal lymphoplasmacytic deciduitis (3)<br>Placental parenchyma: neutrophilic and lymphoplasmacytic placentitis (2), multifocal, global to segmental vascular fibrinoid necrosis with mild neutrophilia (3), and decidua interface necrosis.<br>ISH - Placenta, center cut disc 1: moderate multifocal intense parenchymal staining of scattered placental villi from the chorionic plate to the basal plate, within the outer layers of the chorionic vessels, and within Wharton's jelly of the umbilical cord.<br>ISH - Placenta, center cut disc 2: moderate multifocal intense parenchymal staining of scattered placental villi from the chorionic plate to the basal plate, minimal scattered positive cell staining within the chorionic plate and within chorionic vessels.<br>Umbilical cord: (0).<br>Fetus: diffuse autolysis (3).<br>ISH - Fetus: mild to moderate diffuse positivity in the periosteum, the skeletal muscle of the head, the sclera of the eye, and rare single cell positivity within the neuropil. |
| 91 | II | E | 53 | Uterus: (0).<br>Decidua basalis: multifocal hemosiderosis and multifocal infarction, and rare intralesional vascular fibrinoid necrosis (2).<br>Decidua parietalis: multifocal acute to subacute hemorrhage (3).<br>Placental parenchyma: multifocal, acute villous infarction (3) and multifocal, linear, coagulative necrosis of the decidua-trophoblastic shell interface.<br>ISH - Placenta, center cut disc 1: scattered dense staining within the villous stroma.<br>Fetus: moderate, diffuse, autolysis (3).<br>ISH - Fetus: positive staining of the periosteum of long bones. |
| 92<br>93 | II | F | 175 | Uterus: (0).<br>Decidua basalis: multifocal chronic plasmacytic deciduitis (2).<br>Decidua parietalis: (0).<br>Placental parenchyma: trophoblastic shell-anchoring villi and villi, subacute, focally extensive coagulative and lytic necrosis with multifocal neutrophilic infiltration and rimmed by incipient syncytial trophoblastic cell aggregates (3).<br>ISH Placenta, center cut disc 1: very rare positive cytoplasmic staining of scattered chorionic plate cells.<br>Umbilical cord: (0).<br>Fetus: severe peracute amniotic fluid aspiration, moderate multifocal peracute vascular brain congestion, moderate multifocal thymic hemorrhage with vascular congestion, moderate sinusoid siderophagocytosis and erythrophagocytosis in the mesenteric and axillary lymph nodes, moderate multifocal adrenal corticomedullary and cortical vascular congestion with discrete multifocal hemorrhage, multifocal coalescent testicular vascular congestion (4). |

| | | | | |
|---|---|---|---|---|
| 94 | II | G | 53 | Uterus: (0).<br>Decidua basalis: multifocal neutrophilic deciduitis, chronic thrombosis, and hemosiderosis (1).<br>Decidua parietalis: multifocal neutrophilic deciduitis (1)<br>Placental parenchyma: multifocal degenerative choroid vasculopathy and choroid edema (2), multifocal acute villitis and increased perivillous fibrin (2), multifocal VSVK, focal basal plate thrombosis, and chronic retroplacental hemorrhage (2).<br>ISH - Placenta, center cut disc 1: rare focal parenchymal staining within a single villous.<br>ISH - Placenta, center cut disc 2: scattered minimal single cell positivity within the outer layers of chorionic vessels and the chorionic plate, focal parenchymal staining in scattered single villi.<br>Umbilical cord: degenerative vasculopathy (2).<br>Fetus: diffuse autolysis (3).<br>ISH - Fetus: segmental positive staining of the intestinal serosa, multifocal positive staining throughout cephalic tissues, and positive staining of the periosteum and skeletal muscle of the hind limbs and pelvis.<br>Maternal spleen: diffuse neutrophilic splenitis (2). |
| 95 | II | H | 165 | Natural birth - no maternal/fetal interface tissues obtained |
| 96 | II | I | 165 | Natural birth - no maternal/fetal interface tissues obtained |
| 97<br>98 | II | J | 50 | Uterus: (0).<br>Decidua basalis: minimal multifocal chronic deciduitis (1).<br>Decidua parietalis: (0).<br>Placental parenchyma: focal neutrophilic intervillositis (1).<br>ISH Placenta, center cut disc 1: single cell staining in the wharton’s jelly of the umbilical cord, intense multifocal staining within the villous parenchyma, scattered staining in the chorionic plate, and scattered staining in the outer layers of the large chorionic blood vessels.<br>ISH Placenta, center cut disc 2: intense multifocal villous parenchymal staining, scattered chorionic plate staining, and scattered staining in the outer layers of the large chorionic vessels.<br>Umbilical cord: diffuse hypercoiling (4).<br>Fetus: diffuse autolysis (3), diffuse congestion in the cerebral meninges.<br>Maternal mesenteric lymph node: lymphoid hyperplasia (3).<br>ISH - Fetus: intense focal myocardial staining, intense vertebral and costal (rib) periosteal staining, moderately intense scattered skeletal muscle staining adjacent to the vertebrae, scapula, humerus, pulmonary interstitium, and thoracic pleura, scattered single cell staining in the liver and the renal interstitium, multifocal staining in the sclera with no staining in the retina and lens, and dense staining throughout the nasal turbinates, spinal cord and cerebral meninges, and the cerebral neuropil. |

| | | | | |
|---|---|---|---|---|
| 99 | III | K | 38 | Uterus: (0).<br>Decidua basalis: (0).<br>Decidua parietalis: (0).<br>Placental parenchyma: focally extensive coagulative necrosis with intralesional vascular fibrinoid necrosis (4), and neutrophilia (1).<br>ISH - Placenta, center cut disc 1: marked diffuse villous parenchymal staining from the basal plate to the chorionic trophoblastic shell with transmural segmental sparing of villi.<br>ISH - Placenta, center cut disc 2: marked diffuse villous parenchymal staining from basal plate to trophoblastic shell and mild diffuse scattered chorionic plate staining.<br>Fetus: tissue autolysis (3).<br>ISH - Fetus: marked diffuse nasal turbinate staining, diffuse neuropil staining, and diffuse tissue staining with no osseous (bone) staining. |
| 100 | III | L | 46 | Uterus: multifocal lymphoplasmacytic myometrial vasculitis (3).<br>Decidua basalis: chronic multifocal lymphoplasmacytic and neutrophilic deciduitis, with necrosis and multifocal hemosiderosis (2).<br>Placental parenchyma: multifocal avascular villi (2).<br>ISH- Placenta center cut disc 1: scattered staining of the outer umbilical cord stroma, the chorionic plate, and the chorionic vessels, intense and dense villous parenchymal staining of scattered single placental villi.<br>IISH - Placenta, center cut disc 2: intense and dense villous parenchymal of scattered villi, mild scattered diffuse chorionic plate staining, and outer the layers of the chorionic vessels.<br>Umbilical cord: lymphoplasmacytic vasculitis (2).<br>Fetus: (0).<br>ISH - Fetus: intense dense cellular staining of the nasal turbinates, periosteum, cranial skeletal muscle, the tongue, the dermis of the chin and lower jaw, connective tissue and muscle around the larynx, and the serosal surface of the esophagus and the cerebral neuropil.<br>Maternal mesenteric lymph node: hemosiderosis (3). |
| 101<br>102 | III | M | 46 | Uterus: (0).<br>Decidua basalis: (0).<br>Decidua parietalis: (0).<br>Placental parenchyma: multifocal neutrophilic intervillositis and multifocal VSVK (1), multifocal basal plate necrosis with mild multifocal villous ischemia (2), and minimal multifocal retroplacental hemorrhage.<br>ISH - Placenta, center cut disc 1: minimal staining in outer layers of chorionic vessels, mild diffuse chorionic plate staining, mild multifocal but dense villous parenchymal staining, and mild multifocal basal plate trophoblastic shell staining.<br>ISH - Placenta, center cut disc 2: minimal staining in outer layers of chorionic vessels, minimal to mild diffuse chorionic plate staining, and mild multifocal but dense villous parenchymal staining.<br>Umbilical cord: (0).<br>Fetus: diffuse autolysis (3).<br>ISH - Fetus: positive staining in the persiosteum, intestinal serosa, and neuropil. |

| | | | | |
|---|---|---|---|---|
| 103 | III | N | 46 | Uterus: (0).<br>Decidua basalis: multifocal acute neutrophilic deciduitis (2).<br>Decidua parietalis: (0).<br>Placental parenchyma: focal basal plate trophoblastic hemorrhage (2).<br>ISH - Placenta, center cut disc 1: mild staining in outer layers of chorionic vessels, moderate diffuse staining throughout the chorionic plate, dense villous parenchymal staining of scattered villi.<br>ISH - Placenta, center cut disc 2: mild staining in outer layers of chorionic vessels, moderate diffuse staining throughout the chorionic plate, dense villous parenchymal staining of scattered villi including stem villi.<br>Fetus: diffuse autolysis (4).<br>ISH- Fetus: positive staining in the periosteum and musculature of the head and body tissues.<br>Mesenteric lymph node, maternal: diffuse sinus histiocytosis with minimal hemosiderosis (3). |
| 104 | | | | |

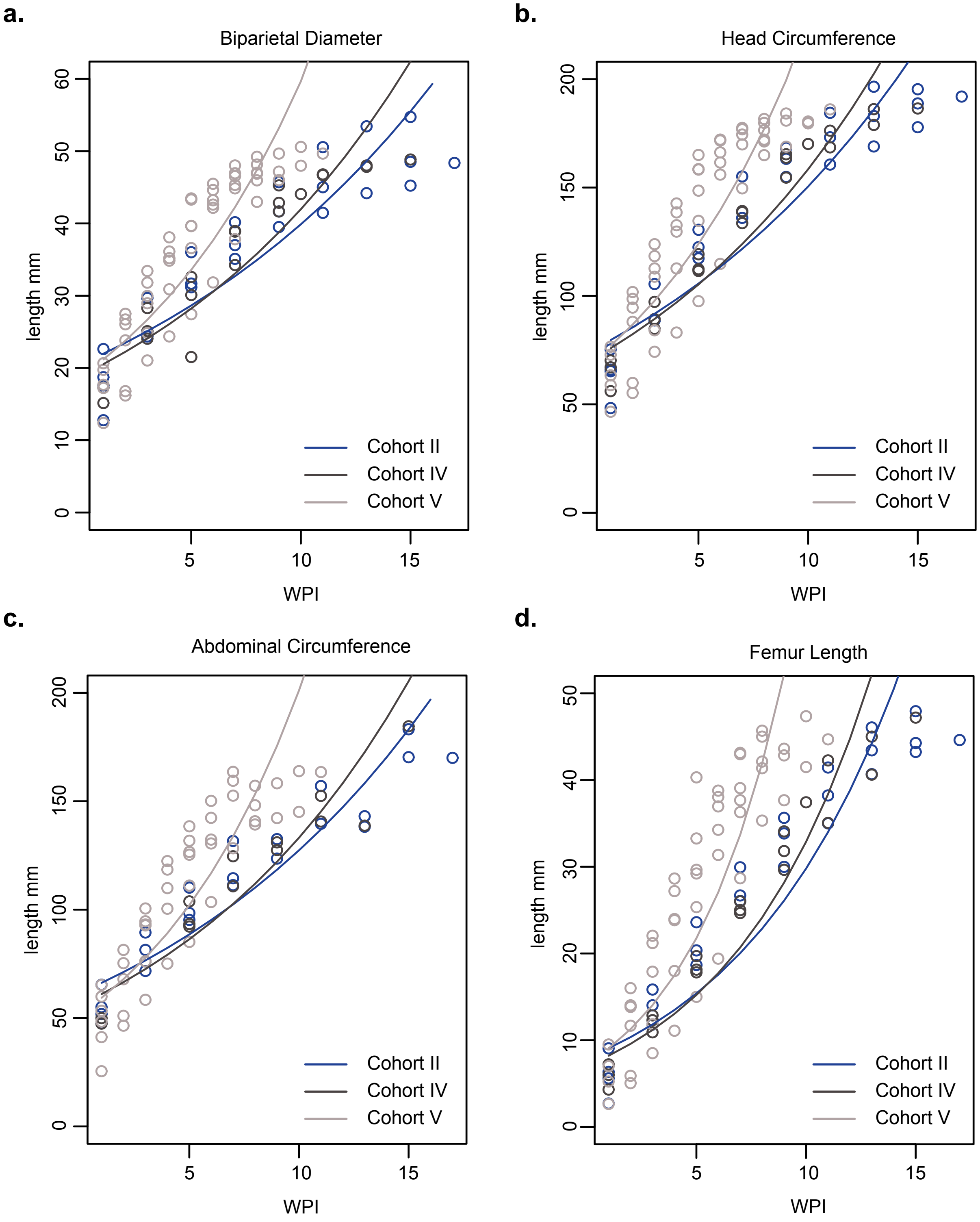


**Supplemental Fig. 3 : In-utero growth trajectories of Abdominal circumference (AC), Biparietal Diameter (BPD), Femur length (FL), and Head circumference (HC) compared between Cohorts II, IV, and V.** Slopes for each Cohort were determined using a linear mixed effects regression model with animal specific measurements and an autoregressive correlation structure over WPI.

**Supplemental Table 5:** Slope estimates (95% CI) for Abdominal circumference (AC), Biparietal Diameter (BPD), Femur length (FL), and Head circumference (HC) on log scale, stratified by cohort. 1: p-value for comparison of slopes between Cohort II vs. Cohort IV; 2: p-value for comparison of slopes between Cohort II vs. Cohort V; 3: p-value for comparison of slopes between Cohort IV vs. Cohort V.

| Outcome | Cohort II | Cohort IV | Cohort V | p-value[1] | p-value[2] | p-value[3] |
|---|---|---|---|---|---|---|
| AC | 0.08 (0.05-0.1) | 0.09 (0.06-0.12) | 0.16 (0.13-0.18) | 0.5809 | 0.0001 | 0.0041 |
| BPD | 0.07 (0.05-0.09) | 0.08 (0.05-0.11) | 0.13 (0.11-0.16) | 0.6396 | 0.0005 | 0.0147 |
| FL | 0.14 (0.1-0.18) | 0.16 (0.1-0.23) | 0.26 (0.21-0.31) | 0.5272 | 0.0005 | 0.0207 |
| HC | 0.07 (0.05-0.1) | 0.08 (0.05-0.11) | 0.13 (0.11-0.16) | 0.6297 | 0.0004 | 0.0120 |

**Supplemental Table 6: Comprehensive ultrasound remarks during pregnancy.** Animals C and M did not have any notable findings.

| Animal ID | Remarks |
|---|---|
| A | On GD 53 (day of pregnancy loss) fetal hydrops, some skin edema, and pleural effusion was observed. Hydrops can be a precursor to fetal demise. |
| B | On GD 44 skin appeared edematous and on GD 47 a thick nuchal fold with lots of skin edema was observed. There were possible septations within the edema consistent with a cystic hygroma. Edematous fetal skin with septations within was observed again on GD 51 and on the day of pregnancy loss (GD 54). In humans, skin edema and cystic hygromas can be seen in cases of aneuploidy, Noonan syndrome, and fetal cardiac anomalies. |
| C | No significant findings. |
| D | On GD 44 debris was present between the amnion-chorion, possibly blood or infection. |
| E | On GD 25 the maternal abdomen and area surrounding the uterus had a lot of free fluid, suggestive of maternal ascites. On the day of pregnancy loss (GD 53) edema in the fetal skin, head, and abdomen was observed along with 0.13cm of fluid surrounding the fetal heart. These findings are consistent with fetal hydrops which can be a precursor to fetal demise. On this day (GD 53) fluid had also collected in the maternal abdomen (ascites) and placental calcifications were observed along the decidual edge. |
| F | Fetal scalp edema was observed on GD 55.Scalp edema is suggestive of fetal hydrops which can be a precursor to fetal demise. One week later (GD 62) 0.17cm fluid was found surrounding the fetal heart. The pericardial fluid volume decreased to 0.07cm by GD 76 and placental calcifications began to appear at the decidual edge. Placental calcifications increased throughout the placenta during pregnancy. |
| G | A 2.63 cm retroplacental bleed was observed between amnion and chorion located over the cervix on GD 36 along with a clot near the fundus which measured 2.32x1.26cm. Similar clots are seen in the setting of placental abruption or occasionally with subchorionic hematomas. Abruption can have significant implications with bleeding from the maternal and fetal circulation and can result in demise of the baby or mother. Subchorionic hematomas often have minimal sequelae and usually resolve. Scant calcifications appeared along the decidual edge on the day of pregnancy loss (GD 53). |
| H | The fetal bowel appeared slightly echogenic on GD 78 and free fluid resembling ascites was observed in the maternal abdomen. On GD 92 the fetal bowel was less echogenic, and ascites were not seen. Placental calcifications began at GD 92 along the decidual edge and increased throughout the placenta during pregnancy. On GD 120 there was mixed echogenicity observed at the pulmonary artery. |

| | | |
|---|---|---|
| 132 | I | On GD 57 the fetal bowel was slightly echogenic and placental calcifications began to appear along the decidual edge. These calcifications increased throughout the placenta during pregnancy. GD 155 1.96cm mass noted in maternal tissues deep to the skin. Fluid in anterior tissue and enlarged pelvic lymph nodes. |
| 133 | J | Copious echogenic debris, possibly blood or infection, between chorion and amnion was first noted on GD 40 and continued until the pregnancy was lost on GD 50. On the day of pregnancy loss, hydrops was observed and impacted the fetal thorax, abdomen (0.28cm fluid), and scalp (0.25cm fluid). |
| 134 | K | On the day of pregnancy loss there was echogenic debris between the amnion and chorion (GD 38). |
| 135 | L | Scant debris between amnion and chorion, possibly blood or infection, was observed on GD 36 and the amnion seemed a little tight around the embryo. This debris increased up until the day of pregnancy loss (GD 46). |
| 136 | M | No significant findings. |
| 137 | N | The day before pregnancy loss (GD 43) there was some swelling in the nuchal and thoracic region of the embryo and echogenic debris was observed between chorion and amnion. The echogenic debris could have been blood or represented infection. The thoracic swelling increased on the day of pregnancy loss (GD 44). Debris and swelling could be related; if blood loss resulted in anemia, then this can result in hydrops, a precursor to fetal demise. |
| 138 | O | Placental calcifications began on GD 76 along the decidual edge and increased throughout the placenta during pregnancy. On GD 118 calcifications were also noted in the right upper lobe of the liver and medial to the stomach. These echogenic foci in the abdomen can be random and unexplained or can be seen in the setting of in utero infection, cystic fibrosis (in humans), intra amniotic bleeding (where the baby swallows blood and this becomes echogenic in the bowel), or genetic conditions like aneuploidy. |
| 139 | P | Placental calcifications began to form along the decidual edge at GD 86 and increased throughout the placenta during pregnancy. On GD 129 the inferior vena cava (IVC) looked large (0.37cm) compared to the aorta (0.25cm). On GD 143 the IVC measured at 0.31cm/0.39cm and appeared to be at the upper limit of normal or slightly dilated. The VC can be enlarged randomly, when the heart isn't pumping as well, as seen in heart failure, or can be seen if there is an aneurysm. |
| 140 | Q | On GD 49 edema was noted around the thorax and abdomen posteriorly measuring 0.16cm. Placental calcifications began to form along the decidual edge at GD 77 and increased throughout the placenta during pregnancy. |
| 141 | R | On GD 55 the fetal bowel is slightly bright but does not appear echogenic. Placental calcifications began at GD 62 along the decidual edge and increased throughout the placenta during pregnancy. |
| 142 | S | On GD 63 the fetal chest wall appeared thick and placental calcifications began to form along the decidual edge. These calcifications increased throughout the placenta during pregnancy. |
| 143 | T | The frontal bones of the fetal calvarium were slightly curved inward on GD 57. Some debris was observed in the fetal stomach on GD 71 and the bowel appeared echogenic. Placental calcifications began at GD 85 along the decidual edge and increased throughout the placenta during pregnancy. There was a mixed echogenicity mass located anteriorly and superior to the heart on GD 127. This mass was primarily hypoechoic but had hyperechoic spots throughout. On GD 141 the cystic mass was anterior to the heart and measured 1.7x 1.34x 0.98 cm. This is indicative of possible CPAM or bronchopulmonary sequestration. On GD 155 the CPAM mass measured: 1.25x 0.97x 0.71cm, volume = 0.45mL, volume ratio = 0.03cmx2. There is a possibility this mass was the thymus. |
| 144 | U | Placental calcifications began to appear along the decidual edge on GD 73 and increased throughout the placenta during pregnancy. |
| 145 | V | Placental calcifications began to appear along the decidual edge on GD 78 and increased throughout the placenta during pregnancy. |
| 146 | M | Placental calcifications began to appear along the decidual edge on GD 68 and increased throughout the placenta during pregnancy. |

147

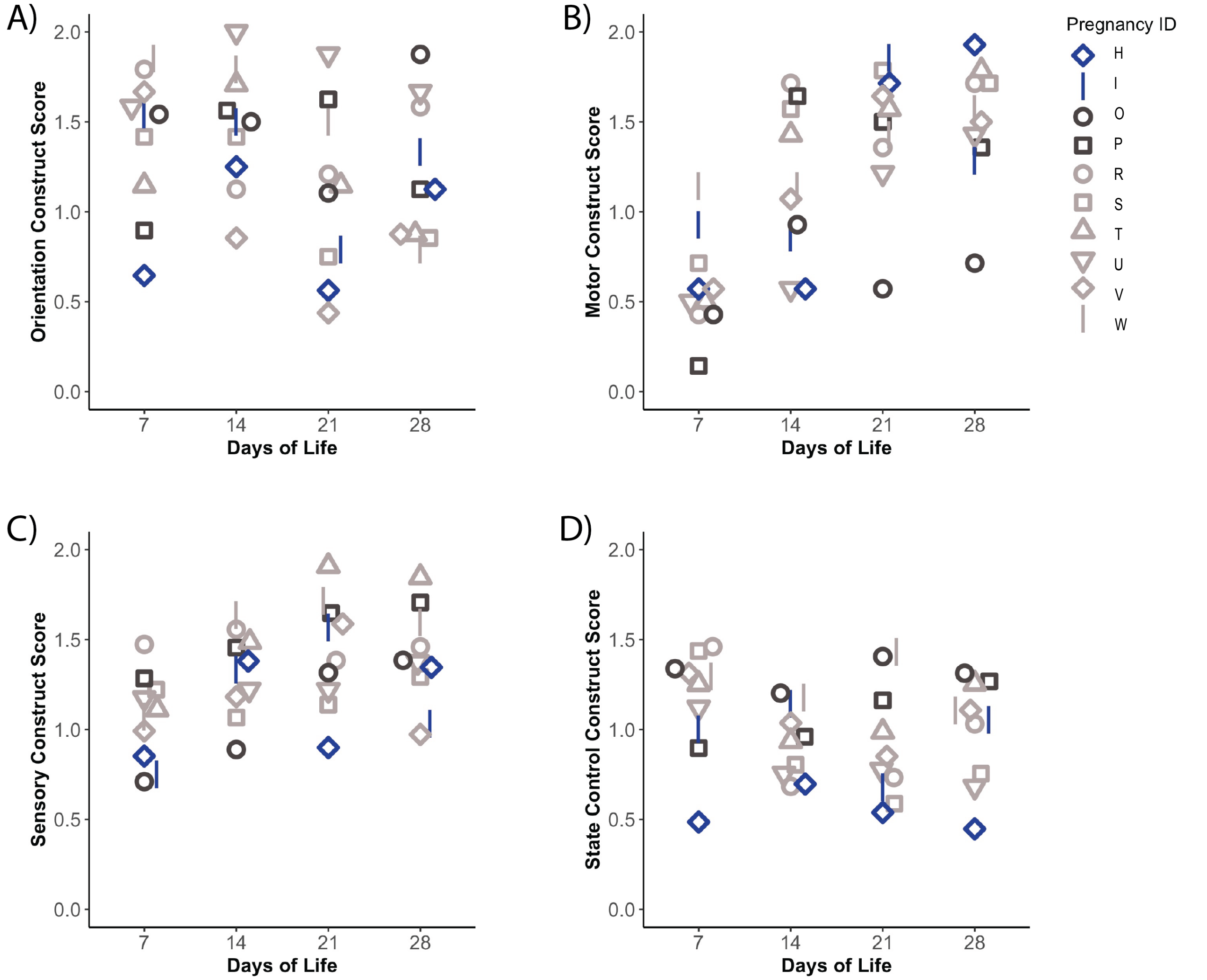


**Supplemental Fig. 4: Neurodevelopmental testing was measured by the SNAP at day 7, 14, 21, and 28 days of life.** The SNAP is composed of 4 main constructs: (a) Orientation, (b) Motor maturity and activity, (c) Sensory responsiveness, and (d) State control. Cohort II (SIV-/ZIKV+ +ART) infants are in blue, Cohort IV (SIV+/ZIKV- +ART) infants are in light gray, and Cohort V (SIV-/ZIKV- +ART) infants are in dark gray. Individual infants are represented by symbols that correspond to their Pregnancy ID.

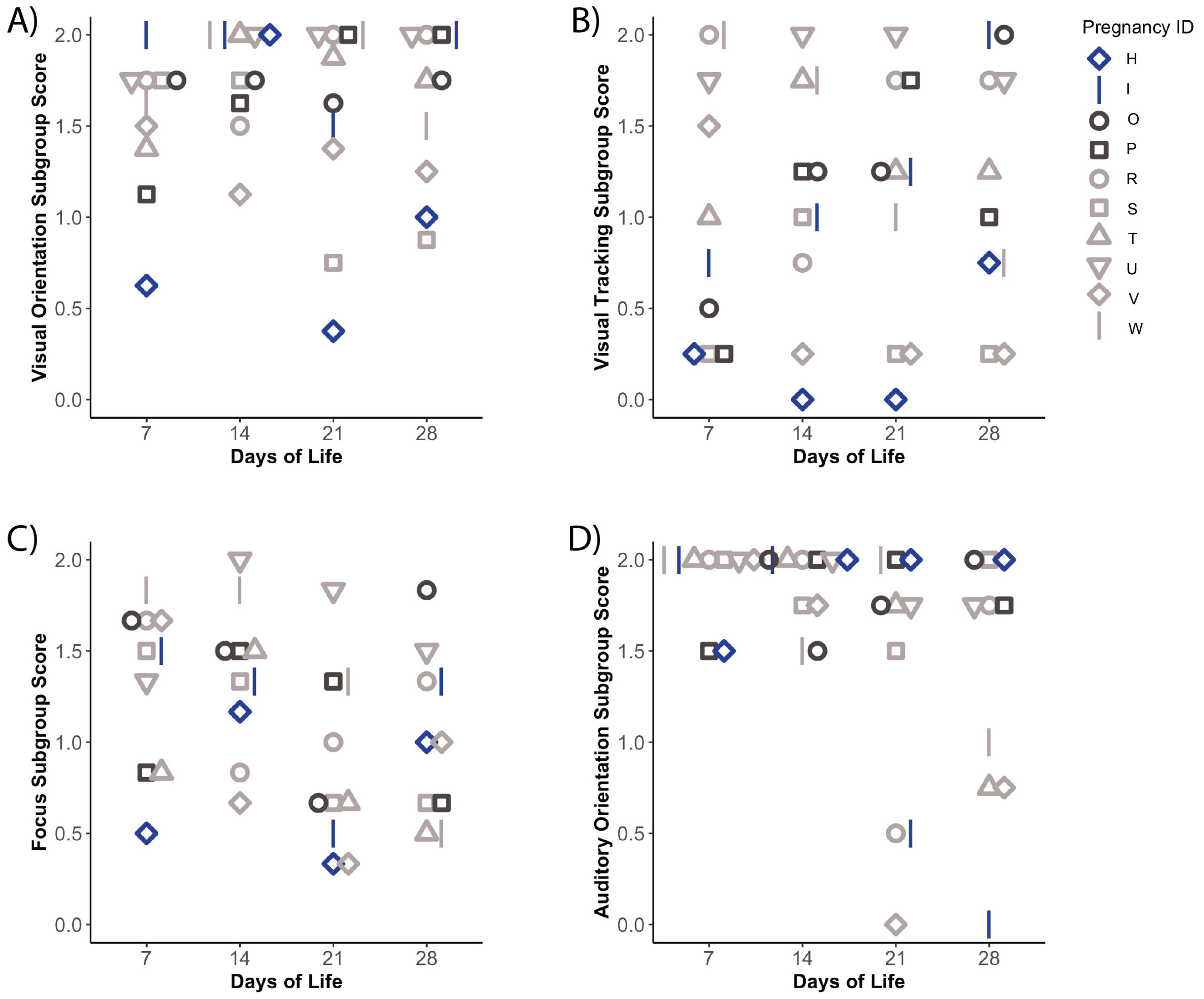


**Supplemental Fig. 5: SNAP Orientation construct was separated by sensory modality and task into 4 main subgroups to see where differences may be occurring.** The SNAP Orientation sub-groups consist of A) Visual orientation, B) Visual tracking, C) Focus, and D) Auditory orientation. Cohort II (SIV-/ZIKV+ +ART) infants are in blue, Cohort IV (SIV+/ZIKV- +ART) infants are in light gray, and Cohort V (SIV-/ZIKV- +ART) infants are in dark gray. Individual infants are represented by symbols that correspond to their Pregnancy ID.

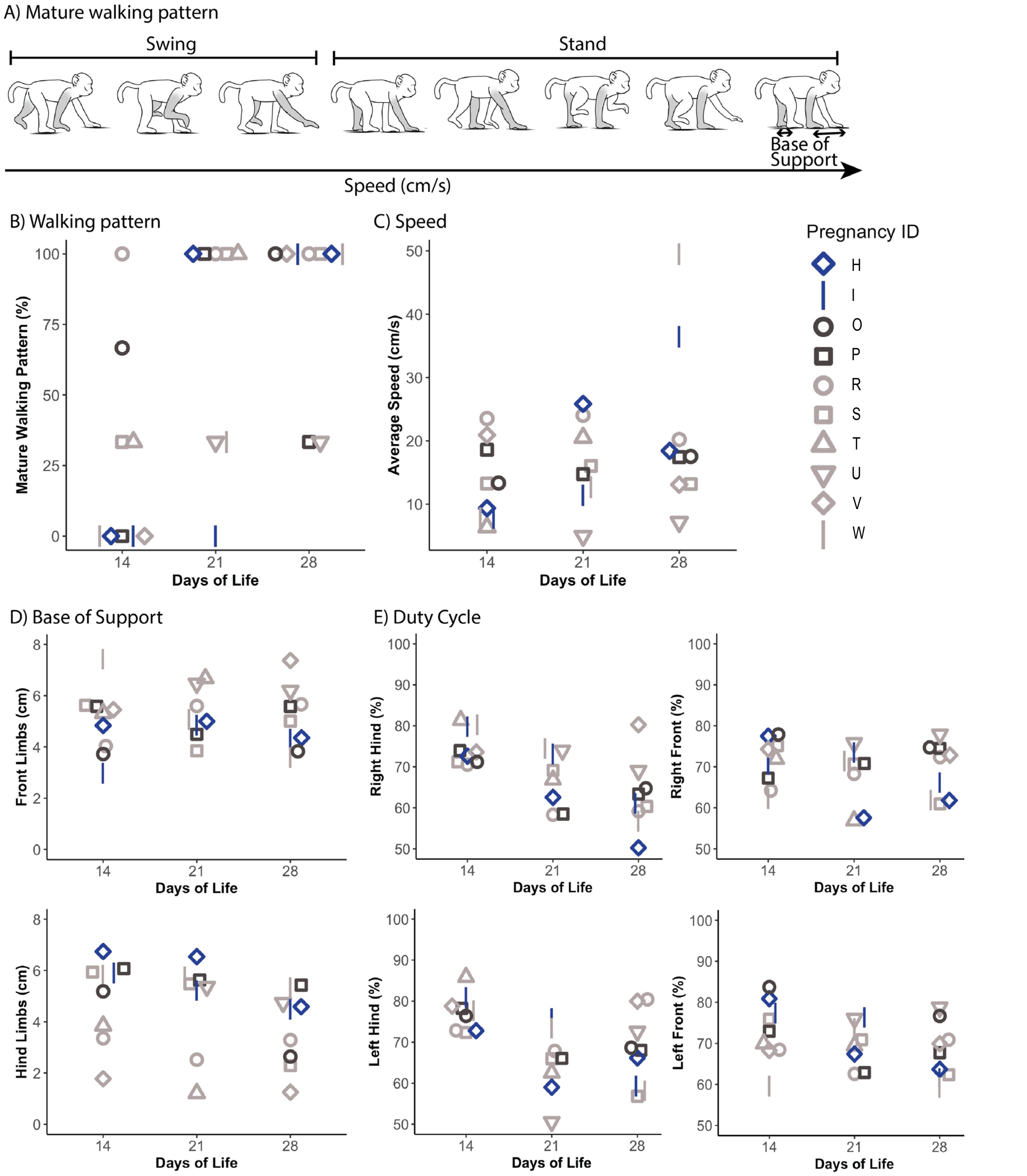


**Supplemental Fig. 6: Infant locomotion was measured at 14, 21, and 28 days of life using the Noldus Catwalk.** (a) Visual representation of the gait variables included mature walking pattern (where the contralateral limbs are moving through the swing and stance phase close together in timing), speed (how fast the infant walked across the catwalk), base of support (distance between right and left limbs), and duty cycle (percent of time the infant is standing on the walkway). (b) Percent of time infants use a mature walking pattern. (c) Average speed across runs, (d) Base of support for the front and hind limbs, (e) Duty cycle time the infant was standing on each limb. Cohort II (SIV-/ZIKV+ +ART) infants are in blue, Cohort IV (SIV+/ZIKV- +ART) infants are in light gray, and Cohort V (SIV-/ZIKV- +ART) infants are in dark gray. Individual infants are represented by symbols that correspond to their Pregnancy ID.

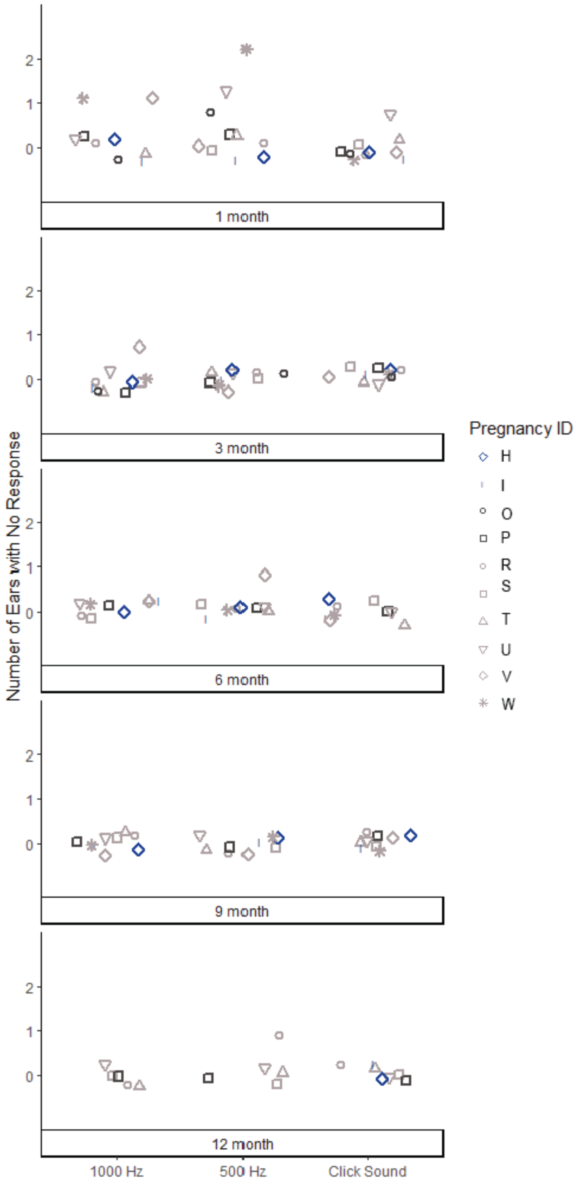


**Supplemental Fig. 7: Hearing: longitudinal auditory brainstem responses.** Infant macaques were tested via auditory brainstem response to click, 1000 Hz, and 500 Hz stimuli. The presence or absence of a Wave IV response for each ear was assessed and summarized for each animal at 1, 3, 6, 9, and 12 months of age. Not all the animals received testing at 12 months of age because some were not 12 months old yet. Infants with normal hearing are expected to have a score of “0” with both ears having a Wave IV auditory brainstem response at the lowest intensity level tested. Cohort II (SIV-/ZIKV+ +ART) infants are in blue, Cohort IV (SIV+/ZIKV- +ART) infants are in light gray, and Cohort V (SIV-/ZIKV- +ART) infants are in dark gray. Individual infants are represented by symbols that correspond to their Pregnancy ID.

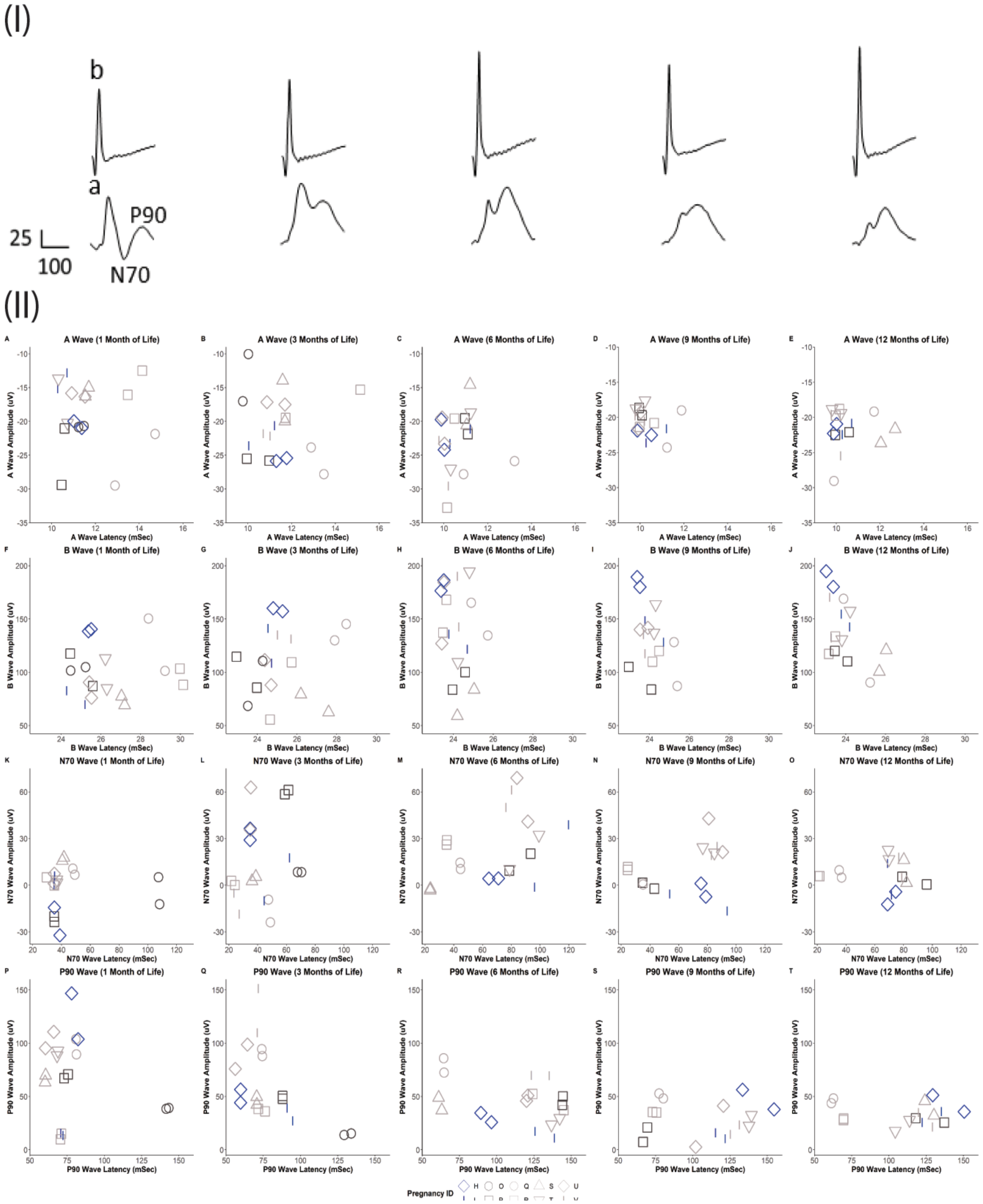


**Supplemental Fig. 8: Visual electrophysiology.** Visual function studies comprising photopic single flash electroretinograms (ERGs) and visual evoked potentials (VEPs). (I) Schematic representing the origins of the A and B waves within an ERG and the N70 and P90 waves within a VEP for each age group, aligned with the ages in part II. These representative waveforms were created from the averages of all the traces for the controls at each time point. (II) ERG A- and B-wave components were measured at 1, 3, 6, 9, and 12 months of age (A-J). The right and left eyes are plotted as individual data points. VEP N70- and P90-wave components were measured at 1, 3, 6, 9, and 12 months of age (K-T). The right and left hemispheres are plotted as individual data points. Cohort II (SIV-/ZIKV+ +ART) infants are in blue, Cohort IV (SIV+/ZIKV- +ART) infants are in light gray, and Cohort V (SIV-/ZIKV- +ART) infants are in dark gray. Individual infants are represented by symbols that correspond to their Pregnancy ID.

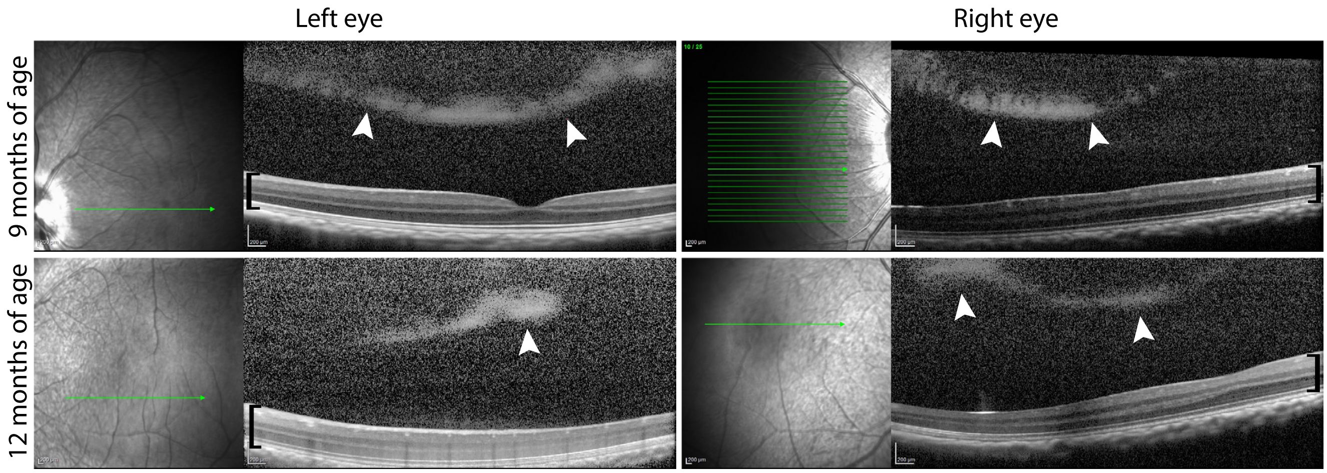


**Supplemental Fig. 9. Retinal and vitreous images of the infant from Pregnancy H.** Optical coherence tomography images of the infant from Pregnancy H at 9 at 12 months of age. Vitreous opacities, or "clumping", is seen both right and left eyes, indicated by the white arrows. The retinal layers are denoted with a black bracket. The green arrow shows the relation of the retinal section shown in relation to the fovea.

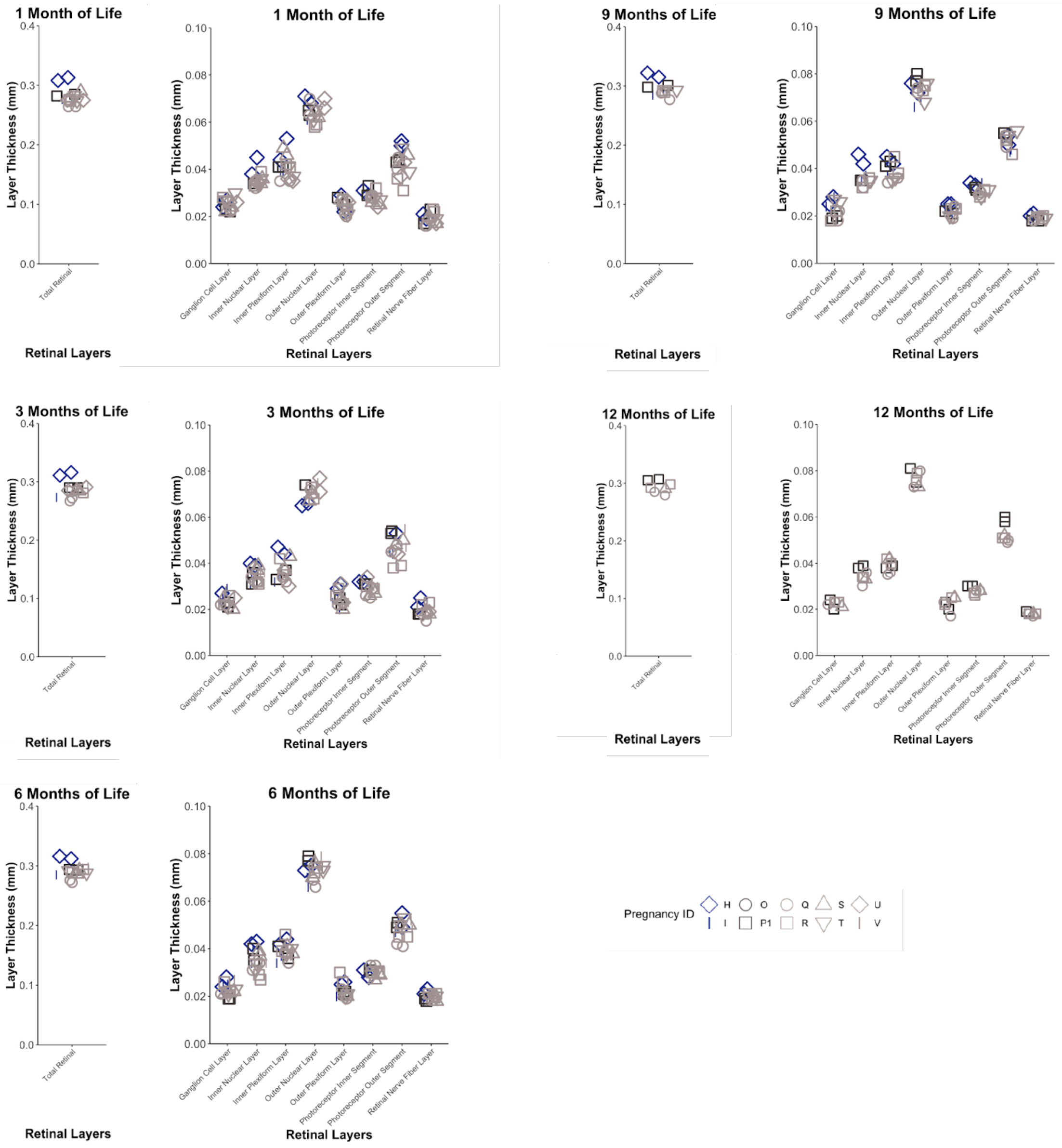


**Supplemental Fig. 10: Optical coherence tomography (OCT): retinal layer thicknesses.** Retinal layer thicknesses were measured by optical coherence tomography (OCT) in infants at 1, 3, 6, 9, and 12 months of age. Total retinal thickness, choroidal thickness, and the thickness of individual retinal layers (ganglion cell layer, inner nuclear layer, inner plexiform layer, outer nuclear layer, outer plexiform layer, photoreceptor inner segment, photoreceptor outer segment, retinal nerve fiber layer) were determined by segmentation. Cohort II (SIV-/ZIKV+ +ART) infants are in blue, Cohort IV (SIV+/ZIKV- +ART) infants are in light gray, and Cohort V (SIV-/ZIKV- +ART) infants are in dark gray. Individual infants are represented by symbols that correspond to their Pregnancy ID.
